# Serpin A3 is Increased in Cerebral Amyloid Angiopathy and ARIA

**DOI:** 10.64898/2026.10.19.759840

**Authors:** Lissa Ventura-Antunes, Vaibhav A. Janve, Neely Womble, Conor J. Weasler, Sahir A. Hai, Hosamulhaq Brula, Bret C. Mobley, Timothy J. Hohman, Wilber Romero-Fernandez, Matthew S. Schrag

## Abstract

Vascular pathology is an important contributor to dementia, and astrocyte dysfunction is increasingly recognized as a key mediator of cerebrovascular injury. Serpin A3 is a well-known marker of reactive astrocytes, yet its role in vascular degeneration in cerebral amyloid angiopathy and Alzheimer’s disease remains poorly defined. We combined analyses of single nucleus transcriptomic datasets with western blotting and quantitative two- and three-dimensional microscopy of human brain tissue to characterize Serpin A3 expression, distribution, cellular sources, and neuropathological correlates in the human brain. Serpin A3 protein abundance was increased in cases with cerebral amyloid angiopathy and Alzheimer’s disease compared to neurological controls, both by western blot and quantitative microscopy. The tissue distribution of Serpin A3 was heterogeneous and was predominantly within astrocytes associated with degenerating arterioles. Serpin A3 was also detected around parenchymal β-amyloid plaques. In a postmortem case of active and severe lecanemab-associated amyloid-related imaging abnormalities, light sheet microscopy revealed prominent Serpin A3 immunoreactivity around inflamed, dilated vessels and microhemorrhages, with Serpin A3 levels higher those found in the Cerebral Amyloid Angiopathy cases. These findings identify Serpin A3 as a marker of cerebrovascular injury and astrocyte-driven inflammatory responses in cerebral amyloid angiopathy.

## Background

Vascular cognitive impairment is the second most common cause of dementia after Alzheimer’s disease (AD) [1]. Although vascular and neurodegenerative pathologies frequently coexist in AD, the mechanistic links underlying their interaction remain incompletely understood.

Astrocytes perform critical roles in maintaining brain homeostasis, including regulating synaptic activity, and participation in the neurovascular unit where it is a central component of the blood-brain barrier (BBB) [2, 3]. Their role in maintaining connective tissue homeostasis and supporting the structural integrity of cerebral vessels is beginning to come into sharper focus [2, 3]. Under pathological conditions, astrocytes undergo reactive phenotypic changes that can promote inflammation and contribute to chronic injury of the cerebral microvasculature.

Increasing evidence identifies reactive astrocytes as important mediators of neurovascular dysfunction in several neurological disorders, including AD and cerebral amyloid angiopathy (CAA). Transcriptomic and spatial profiling approaches have further revealed substantial astrocyte heterogeneity, highlighting distinct subtype-specific anatomical niches and specialized functions relevant to cerebral homeostasis, tissue injury response, and neuropathologies [4, 5].

Our recent publication demonstrated that tumor necrosis factor-α (TNF-α) activates STAT3-associated signaling in cultured astrocytes [6], leading to the secretion of effector molecules including Serpin Family A Member 3 (Serpin A3) that may influence microvascular function. There is growing evidence that the Serpin A3 pathway can also be engaged by additional inflammatory mediators, including IL-1 and IL-6 [7, 8]. Although Serpin A3 has long been recognized as a relatively stringent marker of reactive astrocytes, its mechanistic role in astrocyte biology and cerebrovascular injury remains unclear.

Serpin A3, historically referred to as α1-antichymotrypsin (ACT), is a secreted, glycosylated serine protease inhibitor expressed in numerous tissues, including the central nervous system. Extensive glycosylation contributes to its slower electrophoretic migration relative to its predicted native molecular weight of approximately 47 kDa [9]. Serpin A3 either covalently binds to its target protease or else is cleaved by its target protease and is thus consumed in the process of interacting with its protease [10–12]. Accordingly, Serpin A3 levels are relatively low in normal brain tissue but may become elevated under inflammatory and injury-associated conditions. Serpin A3n, the murine homolog of human Serpin A3, was recently shown to be a marker of injury responsive astrocytes in a murine stroke model [13].

Serpin A3 is reportedly increased in AD brain and in prion diseases [14] as well as in mouse models of AD [15], but how it associates with vascular disease and vascular injury including CAA is not well known. It is noteworthy that CAA has long been associated with a SNP in ACT, which encodes Serpin A3 [16].

Given the potential role of Serpin A3 in vascular remodeling and/or regulating barrier properties, in this study we confirmed its protein level alterations and characterized its distribution in the human brain using quantitative two-and three-dimensional microscopy. We assessed its association with CAA, vascular degeneration, and inflammatory vascular changes associated with amyloid-related imaging abnormalities (ARIA).

## Methods

### Human brain tissue

Brain tissue was obtained from the Vanderbilt Brain and Biospecimen Bank at Vanderbilt University Medical Center, Nashville, Tennessee, USA (IRB # 180287). Written informed consent for each anatomical donation was obtained from the patient or their surrogate decision-maker. The Vanderbilt University Medical Center Institutional Review Board provided ethical supervision, and the study was performed in accordance with the ethical principles of the Declaration of Helsinki. See the supplementary table 1 with neuropathological information for each brain donor.

For donors with CAA, clinical neuroimaging confirmed microhemorrhages in a typical lobar pattern via the Boston criteria, and routine clinical autopsies confirmed and graded the presence of CAA via the Vonsattel criteria.

Blocks of postmortem human brain tissue were selected from the frontal lobe, along with tissue from any anatomical area with microhemorrhages visible upon gross inspection. This sampling approach ensured both a consistent anatomical site and the presence of degenerating vessels. We dissected tissue blocks measuring 1-3 cm^3^ in volume for tissue clearing, and adjacent regions from the same anatomical areas were processed for conventional sectioning at 50 µm on a vibratome for immunohistochemistry. Freshly acquired tissue was fixed in 4% paraformaldehyde (PFA) for 3-5 days prior to clearing and sectioning.

### Antibodies and reagents

Primary antibodies used in this study include rabbit anti-Serpin A3 (GTX55503, GeneTex, CA), rat anti-GFAP (ab279291, Abcam, Cambridge, UK), mouse anti-Actin (66009-1, Proteintech, Rosemont, IL), and mouse anti-Collagen IV (H4B4, DSHB, Iowa, IA).

Secondary antibodies used in this study include anti-rabbit IgG (H+L)-Alexa Fluor 488 (Abcam), anti-rat IgG (H+L) 594 (Abcam), anti-mouse IgG (H+L) 647 (Abcam), anti-rabbit IgG (H+L)-HRP (65-6120; Invitrogen, Carlsbad, CA), IRDye® 680CW donkey anti-mouse (926-68072; LI-COR).

Reagents used in this study include Tomato-Lectin Alexa Fluor 647 (Lectin), 1 µM 4,4’-[(2-methoxy-1,4-phenylene) di-(1*E*)-2,1-ethenediyl] bisphenol (MX-04) (Tocris, Minneapolis, MN), 4 ’, 6-diamidino-2-phenylindol (1 µM, DAPI, Cayman Chemical, Ann Arbor, MI). Purified human Serpin A3 recombinant protein (abx260557, abbexa, Houston, TX). Human Serpin A3 ORF mammalian expression plasmid with C-GFPSpark tag (HG10307-ACG, Sino Biological Inc., Houston, TX). Information on all other reagents is provided in the corresponding sections.

### Two-dimensional immunostaining

Human brain tissue was prepared as previously described [17]. Immunohistochemical labeling was performed as described [18–20]. Free-floating sections were immunolabeled with antibodies against Serpin A3 (10 µg/mL), GFAP (10 µg/mL), and Collagen-IV (10 µg/mL) at 4°C for 3 days. Following extensive washes, sections were incubated with corresponding secondary antibodies (1:1000) for 2 hours, and subsequently counterstaining with MX-04 (1:1000) at room temperature for 30 minutes. Tissues were extensively washed, mounted and imaged using a Zeiss LSM7 confocal microscope (Carl Zeiss Microscopy GmbH, Germany). Mean fluorescence intensity in hippocampal images was quantified using Fiji software and background-corrected by subtracting the average fluorescence intensity of multiple non-staining background regions from the corresponding overall image intensity measurement.

### Transcriptomic analyses from existing datasets

Human single-nucleus transcriptomic analysis utilized the post-quality control (post-QC) dataset consisting of 424 ROS/MAP participants collected by the Rush ADRC and processed at Columbia University Medical Center (AD Knowledge Portal Accession Number: syn31512863). The mean age at death was 89 years, 68% were females, and 37% were AD cases.

Detailed processing methods are described in Fujita et al. 2024 [21]. Briefly following sample processing, library preparation and sequencing of single nuclei, processing of snRNA-seq reads, quality control, cell type classification, and cell subclass analysis were performed. The final dataset contained 300 unique participants with a median of 3824 sequenced nuclei and 1.64 million nuclei in total. All cell type clusters were adapted from the original publication. Eight major cell types including astrocytes, vascular niches (referred to as “endothelial cells” in this article), microglia, oligodendrocytes, oligodendrocyte precursor cells (OPCs), inhibitory neurons, and CUX2+ and CUX2-excitatory neurons. We pooled inhibitory neurons, CUX2+, and CUX2–classes into neurons; oligodendrocytes, and oligodendrocyte precursor cells (OPCs) into oligodendrocytes. In addition, the original cell type subgroups in vascular niches were also retained. Genes with expression in at least 10% of all cells were retained. Cells were removed if they had over 20,000 or less than 200 total RNA with unique molecular identifiers (UMIs) or had over 5% mitochondrially mapped reads. RNA molecular count data were normalized and scaled by the sctransform R package 44 (version 2) using “percent.mt” values for “vars.to.regress” parameter. Additionally, we re-analyzed a publicly available dataset produced by Siciliano et al 2025 [22].

### Cell culture and cDNA transfection

HeLa cells (CCL-2^TM^, ATCC) were grown in DMEM (Gibco) supplemented with 10% (v/v) fetal bovine serum (FBS, Corning), 100 units/mL of penicillin and 100 µg/mL streptomycin (Thermo Fisher Scientific) at 37°C in a humidified atmosphere of 5% CO_2_. For plasmid transfection, plasmid DNA was transfected into cells with Lipofectamine 3000 reagent following the manufacturer’s protocol (L3000015, Invitrogen). The cells were then fixed following immunocytochemistry as described above. Subcellular fluorescence was detected using a Zeiss LSM7 confocal microscope.

### Lightsheet microscopy and three-dimensional immunohistochemistry methods

Optical tissue clearing was performed using previously established protocols [23].

Cleared tissue blocks were immunolabeled with antibodies against Serpin A3 (5 µg/mL) and GFAP (5 µg/mL) at 37°C for 2-3 days. Following an overnight rinse in PBST, samples were incubated with corresponding secondary antibodies (1:500), followed by overnight counterstaining with Tomato-Lectin Alexa Fluor 647 (5 µg/mL) and MX-04 (1:1000). Prior to imaging, tissues were incubated overnight in 63% thiodiethanol and subsequently imaged using a Light Sheet Fluorescence Microscopy (LSFM, Vanderbilt CISR).

### Imaging processes and analyses

Three-dimensional light-sheet image datasets were analyzed using Imaris software (version 11, Bitplane Oxford Instruments). Surface rendering was performed for each structure to visualize the spatial distribution of Serpin A3, GFAP, β-amyloid, and vasculature. The vascular surface rendering obtained from lectin staining excluded microglial cells (which are also stained with Tomato-Lectin). Using a machine learning segmentation tool from IMARIS, which defines objects associated with the vasculature and excludes background and cells not associated with vascular segmentation. Spatial overlap between Serpin A3 and the other structures was then assessed to identify Serpin A3^+^ association with vessels, astrocytes and β-amyloid. For two-dimensional confocal images, fluorescence intensity of each field was quantified across the entire image using Zeiss ZEN software. Colocalization analysis was performed in ZEN by comparing the fluorescence distribution of Serpin A3 with the β-amyloid, GFAP, and vessels on a pixel-by-pixel basis within the same image, enabling assessment of signal overlap between channels. Degenerating arterioles were defined by enlargement, loss of organization of β-amyloid deposition, and irregularity of the vessel wall, consistent with our published criteria [23].

### Protein extraction, SDS-PAGE, and western blot

Cell protein homogenates from grey matter tissue lysates were prepared as described previously [18, 20]. SDS-PAGE and western blot were performed as described previously [18, 20].

### Statistical analysis

The number of samples (*n*) in each experimental group is indicated in the corresponding figure legends. For statistical purposes, reported *n* reflects the number of individual human donors unless explicitly stated. Statistical analyses and graph generation were performed using GraphPad Prism (version 10.0.2, GraphPad Software, Inc., San Diego, CA) a. Comparisons between two groups were performed Welch’s unequal variances *t*-test, or Mann-Whitney U test, as appropriate.

Statistical comparisons in human single-nucleus transcriptomic analysis were performed using Pseudobulk Welch’s t-test on SCTransform-normalized single-cell expression values. Tests were conducted separately within each vascular cell type across phenotype groups. Comparisons were excluded only when both groups exhibited zero variance or fewer than two cells. Welch’s t-test was specifically selected for the patient-level aggregated data to robustly accommodate potential unequal variances and unequal sample sizes between the phenotype groups.

For visualization of subject-level correlations, normalized expression values (SCTransform ’data’ slot) were extracted and averaged across all cells of a given cell type for each individual (pseudobulk mean). This approach controls variance in cell capture and sequencing depth across donors, providing a standardized metric of patient-level cell-type-specific expression to plot against longitudinal cognition scores.

The statistical test used for each analysis is indicated in the corresponding figure legends.

Statistical significance threshold was set *a priori* to the Benjamini-Hochberg method FDR-adjusted *p*-values < 0.05.

## Results

### Serpin A3 protein is increased in Cerebral Amyloid Angiopathy and Alzheimer’s disease

We sought to define the relationship between Serpin A3 and vascular pathology by measuring its level, distribution, and cellular sources in the human brain in the context of CAA, a major vasculopathy associated with AD.

To assess the cell types which express Serpin A3 in human brain, we turned to transcriptomic analyses of the ROSMAP cohort showed that Serpin A3 expression was most expressed in astrocytes, in the vascular compartment (particularly in perivascular fibroblasts cells and endothelial cells) with lower levels of expression in oligodendrocytes, neurons, and other cell types (Fig. 1A and 1B) within the dorsolateral prefrontal cortex. Serpin A3 transcript levels were significantly reduced in AD/CAA only in the vascular compartment, although when stratified by specific cellular subtypes, none reached statistical significance (Fig. 1A and 1B).

**Figure 1.**
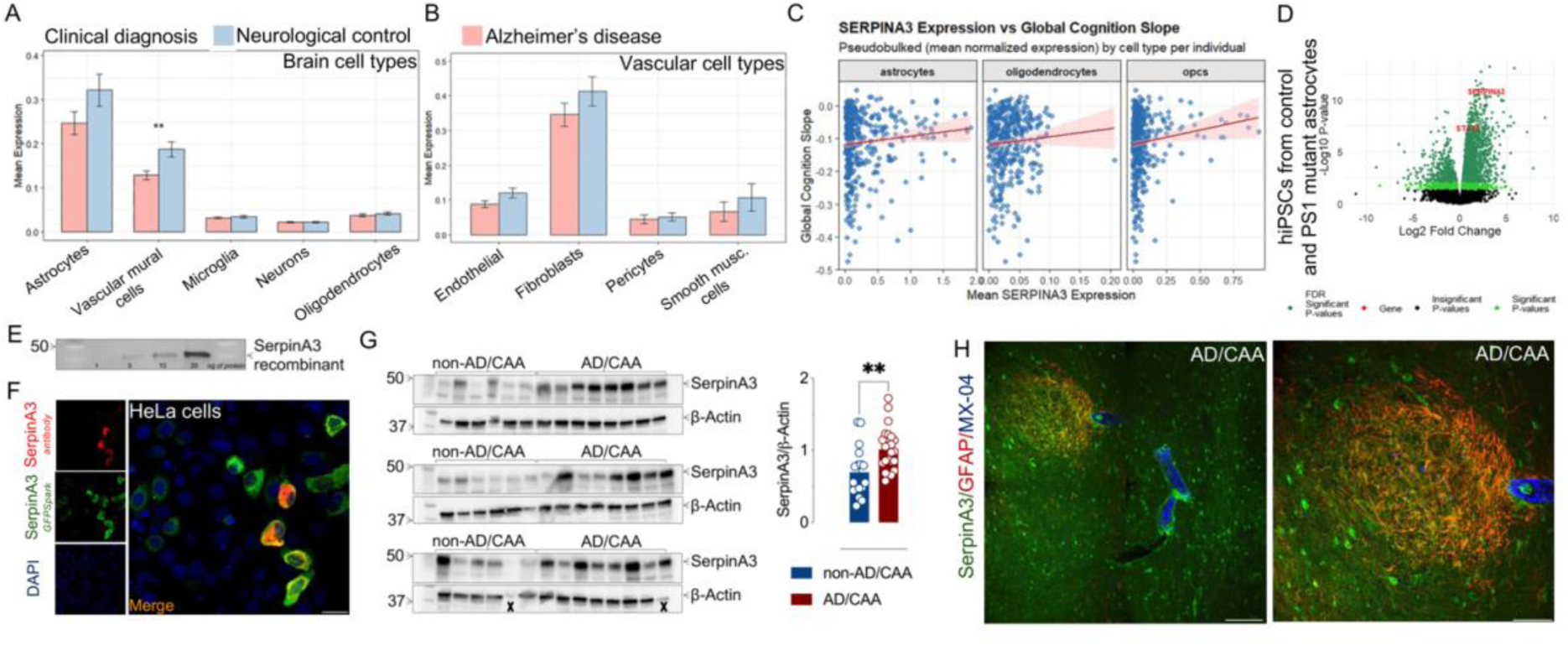
Serpin A3 protein is increased in Cerebral Amyloid Angiopathy and Alzheimer’s disease. A) Transcriptomic analyses of Serpin A3 expression across brain cell types in the ROSMAP cohort. Astrocytes (143,425 cells), vascular mural cells (10,967 cells), microglia (53,969 cells), neurons (146,921 cells), and oligodendrocytes (249,345 cells) obtained from *n*=157 AD/CAA and *n*=143 non-AD/CAA. \*\**p*<0.01 by Welch’s unequal variances *t*-test. B) Transcriptomic analyses of Serpin A3 expression across vascular mural cell subtypes in the ROSMAP cohort. Endothelial cells (5,868 cells from *n*=157 AD/CAA and *n*=143 non-AD/CAA), fibroblasts (2,276 cells from *n*=150 AD/CAA and *n*=140 non-AD/CAA), pericytes (1,825 cells from *n*=152 AD/CAA and *n*=138 non-AD/CAA), and smooth muscle cells (598 cells from *n*=86 AD/CAA and *n*=88 non-AD/CAA). There were no detectable differences between non-AD/CAA and AD/CAA based on Welch’s unequal variances *t*-test. C) Association between SERPIN A3 expression and cognitive decline. Pseudobulk mean SERPIN A3 expression per individual was plotted against global cognition slope across astrocytes, oligodendrocytes, and OPCs. Lines indicate linear regression fits with 95% confidence intervals. D) Volcano plot analysis of differential gene expression in hiPSC-derived astrocytes carrying PS1 mutations relative to control astrocytes (re-analysis of publicly available dataset produced by Siciliano et al., 2025 [22]). Significantly modulated genes are shown in green, and non-significant genes are shown in black. Among the upregulated genes, Serpin A3 and STAT3 were highlighted in red. Statistical significance was determined by false discovery rate (FDR)-adjusted *p* values. E) Validation of the rabbit anti-Serpin A3 antibody for western blotting. Increasing amounts of purified Serpin A3 were probed by western blot and detected a single, specific immunoreactive band around 50 kDa (*n*=3). F) Validation of the rabbit anti-Serpin A3 antibody for immunohistochemistry. HeLa cells transiently expressing fluorescently tagged Serpin A3 (green) were immunolabeled using the anti-Serpin A3 antibody (red), where the tagged protein and antibody signal showed overlapping distributions. Nuclei were counterstained with DAPI (blue), *n*=3. Scale bar, 50 µm. G) Western blot analysis of Serpin A3 protein levels in grey matter tissue from non-AD/CAA (*n*=17) and AD/CAA brains (*n*=23), with ß-actin as a loading control. Quantification (Serpin A3/ ß-actin ratio) is presented as mean ± S.E.M. *\*\*p*<0.01 by using Welch’s unequal variances *t*-test. X; excluded samples. H) Representative confocal images of Serpin A3 immunoreactivity (green) in grey matter of AD/CAA brains obtained using the validated anti-Serpin A3 antibody (*n*=6). Astrocytes are labeled with GFAP (red), β-amyloid deposits, neuritic plaques, neurofibrillary tangles, and other tau aggregates with Methoxy-04 (MX-04, blue). Inset shows a higher-magnification view. Scale bars, 200 µm (main image) and 100 µm (inset).

The general trend toward decreased Serpin A3 expression contrasts previous reports of increased Serpin A3 protein abundance in AD cases [24], although mRNA and protein measures can certainly be discordant. Astrocytic and oligodendrocyte precursor cell expression Serpin A3 correlated with longitudinal cognitive measures; higher levels of expression correlated with better cognitive performance. To further examine astrocytic Serpin A3 regulation in AD-related contexts, we re-analyzed a published RNA-seq dataset from cultured iPSC-derived astrocytes carrying AD-associated presenilin 1 (PSEN1) mutations [22]. In this model, Serpin A3 and STAT3 expression were both increased relative to controls (Fig. 1D).

Next, we quantified Serpin A3 protein abundance in human brains from AD/CAA cases and non-AD/CAA neurological control cases by western blot and immunohistochemistry.

Antibody specificity was validated by detection of a single band corresponding to purified recombinant Serpin A3 by western blot (Fig. 1E and Fig. S1A) and by demonstrating the antibody recognition of fluorescence patterns from transiently expressed GFPS-labeled Serpin A3 (Fig. 1F and Fig. S1B). As previously established, native Serpin A3 from human tissue migrated to a slightly heavier molecular weight than the recombinant form due to glycosylation [9]. As a positive histological control, we stained in grey matter from AD/CAA brains, selecting regions with focal gliosis corresponding to old microhemorrhages or microinfarcts. Serpin A3 is known to be expressed in injury-responsive astrocytes, so we expected robust Serpin A3 expression in these sites.

Serpin A3 protein abundance was significantly increased in posterior frontal cortex AD/CAA relative to non-AD/CAA based on western blot of brain lysates from 23 AD/CAA cases and 17 non-AD/CAA cases (Fig. 1G and Fig. S2). Serpin A3 was also quantified from immunofluorescence of tissue sections and three-dimensional microscopy of optically cleared tissue blocks (Fig. 3 and 4). Immunoreactivity in thin sections from the posterior frontal lobe showed a predominantly vascular and astrocyte-associated distribution, in agreement with the snRNA-Seq data, and was particularly enriched in regions of degenerative vascular pathology (Fig. 1H), including CAA-affected vessels with microaneurysmal changes (Fig 3F). There was minimal Serpin A3 staining of capillaries.

Serpin A3 immunoreactivity was also increased in the hippocampus of AD/CAA cases compared with non-AD/CAA cases (Fig. 2). This increase was evident across the examined hippocampal regions and was characterized by a heterogeneous distribution. Serpin A3 again associated with astrocytic and vascular structures, including some capillaries. In some cases we observed immunoreactivity in cells with neuronal-like morphology. Quantitative analysis confirmed significantly higher Serpin A3 levels in AD/CAA (Fig. 2E).

**Figure 2.**
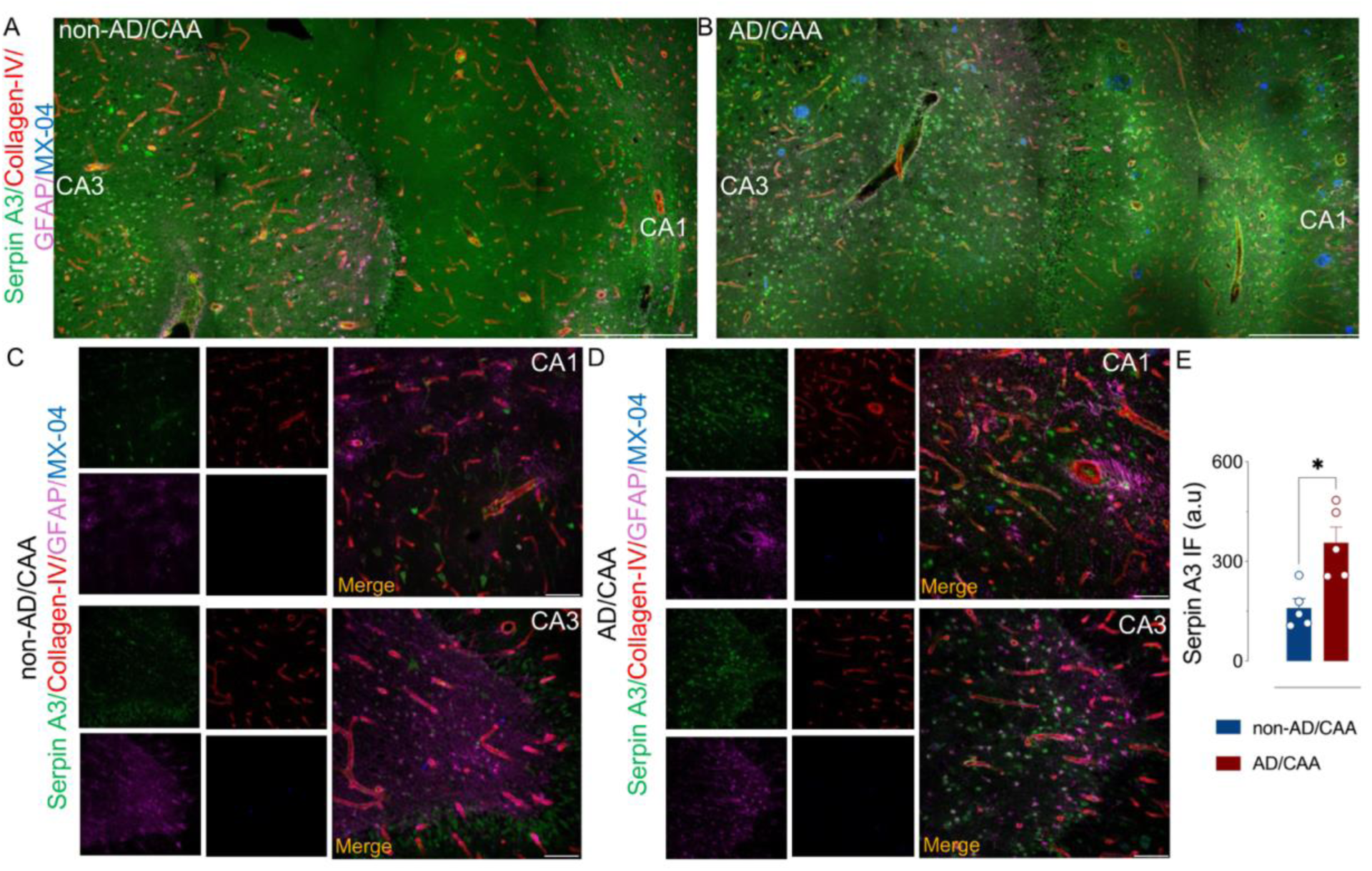
Serpin A3 protein is increased in human hippocampus in Cerebral Amyloid Angiopathy and Alzheimer’s disease. A-B) Representative confocal tile-scan images showing a large area of the human hippocampus from non-AD/CAA (A) and AD/CAA (B) cases. Serpin A3 (green), Collagen-IV (red), GFAP (magenta), and MX-04 (blue). Scale bars, 500 µm C-D) Representative higher magnification images of the CA1 and CA3 regions from non-AD/CAA (C) and AD/CAA (D) cases. Serpin A3 (green), Collagen-IV (red), GFAP (magenta), and MX-04 (blue). Scale bars, 100 µm E) Quantification of Serpin A3 immunofluorescence in non-AD/CAA and AD/CAA hippocampal tissue. Three images were acquired per donor and averaged to obtain one donor-level value. Data are presented as mean ± S.E.M., *n*=5 donors per group \**p*<0.05 by Mann-Whitney U test.

### Serpin A3 is present around degenerating arterioles in cerebral amyloid angiopathy

We compared and quantified histological staining in tissue frontal lobe tissue sections between non-AD/CAA and AD/CAA, which was consistent with the western blot result. We observed a significant increase in Serpin A3 immunoreactivity in AD/CAA cases relative to non-AD/CAA, especially around degenerating vessels (Fig. 3). Serpin A3 levels displayed notable inter-case heterogeneity within the AD/CAA group (Fig. 3G). In non-AD/CAA cases, Serpin A3 immunoreactivity was generally low (Fig. 3A). Occasional non-AD/CAA cases lacking β-amyloid deposition, but with a history of vascular disease showed detectable Serpin A3 immunoreactivity in both the vasculature and parenchyma, resembling the pattern observed in AD/CAA cases (Fig. 3B). In AD/CAA cases, Serpin A3 upregulation was predominantly associated with degenerating arterioles (Fig. 3C-F). This was particularly evident in vessels exhibiting microaneurysmal changes, onion-skinning, rupture, and/or heavy amyloid deposition with distortion of the underlying vascular anatomy. Marked Serpin A3 accumulation was present within degenerating vessel walls and extending into the adjacent parenchyma (Fig. 3D). The statistical correlation between Serpin A3 and a vascular marker or β-amyloid staining was similar in AD/CAA and non-AD/CAA cases (see Fig. 3B). To further clarify the relationship between these structures, we conducted analyses in optically cleared three-dimensional tissue blocks, which we hypothesized would better preserve the relationship between these complex structures.

**Figure 3.**
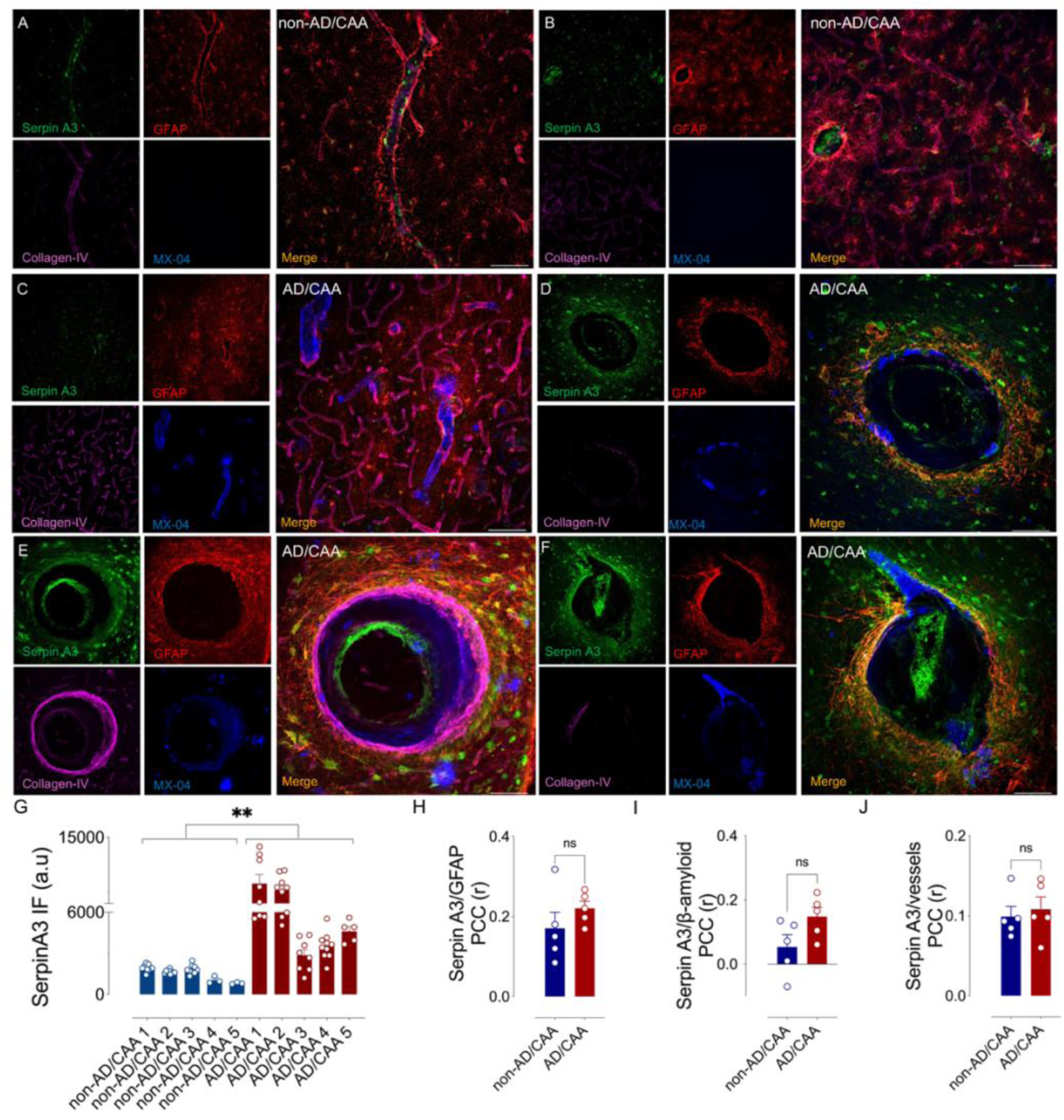
Serpin A3 was increased in Alzheimer’s disease and cerebral amyloid angiopathy in posterior frontal lobe brain tissue. A-F) Representative confocal images of Serpin A3 immunoreactivity (green) in tissue sections of human cortex. Astrocytes are labeled with GFAP (red), vascular basement membrane with Collagen-IV (magenta), and β-amyloid deposits, neuritic plaques, neurofibrillary tangles, and other tau aggregates with Methoxy-04 (MX-04, blue). Scale bars, 100 µm. A and B) Non-AD/CAA cases have minor perivascular staining of Serpin A3. C) Serpin A3 is similarly present at low levels in the microvasculature in AD/CAA in the presence of β-amyloid deposition when the arteriole is structurally intact. D) Serpin A3 level is markedly induced around vessels with degenerating features, including enlarged diameter and fragmented β-amyloid deposits. E and F) Serpin A3 level becomes markedly increased as the severity of degeneration increases. (E) An arteriole with thickened vessel wall and concentric “onion skinning” or “vessel within a vessel” morphology and (F) a vessel showing microaneurysmal degeneration, which is a precursor to rupture, prominently associate with Serpin A3 in GFAP-positive reactive astrocytes. G) Quantification of Serpin A3 immunofluorescence in thin sections of posterior frontal lobe tissue. Bars show all measurements for each donor, illustrating heterogeneity within and between groups. For statistical analysis, measurements were averaged for each patient, and each average was treated as one independent biological replicate (*n*=5 patients per group). Data are presented as mean ± S.E.M. \*\**p*<0.01 by Mann-Whitney U test. H-J) Pearson correlation coefficient (PCC, r) measuring the association between (H) Serpin A3 and GFAP; (I) Serpin A3 and β-amyloid and (J) Serpin A3 and Vessels. Data are presented as mean ± S.E.M., from *n*=5 donors per group. No differences between groups were detected using Mann-Whitney U test

### Serpin A3 was associated with ß-amyloid, GFAP-positive astrocytes, and cerebral amyloid angiopathy in three-dimensional images

We performed three-dimensional image analysis using images obtained from optically cleared tissue blocks and light-sheet microscopy. We generated volume renderings to quantitatively assess the spatial association of Serpin A3 with astrocytes, β-amyloid, and blood vessels. We found that in AD/CAA cases there was a spatial association of Serpin A3 with GFAP positive astrocytes and β-amyloid plaques, in addition to the association with degenerating vessels (Fig. 4A). Volume rendered reconstructions demonstrated higher levels of Serpin A3 in GFAP-positive astrocytes in AD/CAA cases compared with non-AD/CAA cases (Fig. 4C).

Additionally, we found that Serpin A3 was more abundant in β-amyloid deposits and in vessels in cases with CAA and AD (Fig. 4D and E).

**Figure 4.**
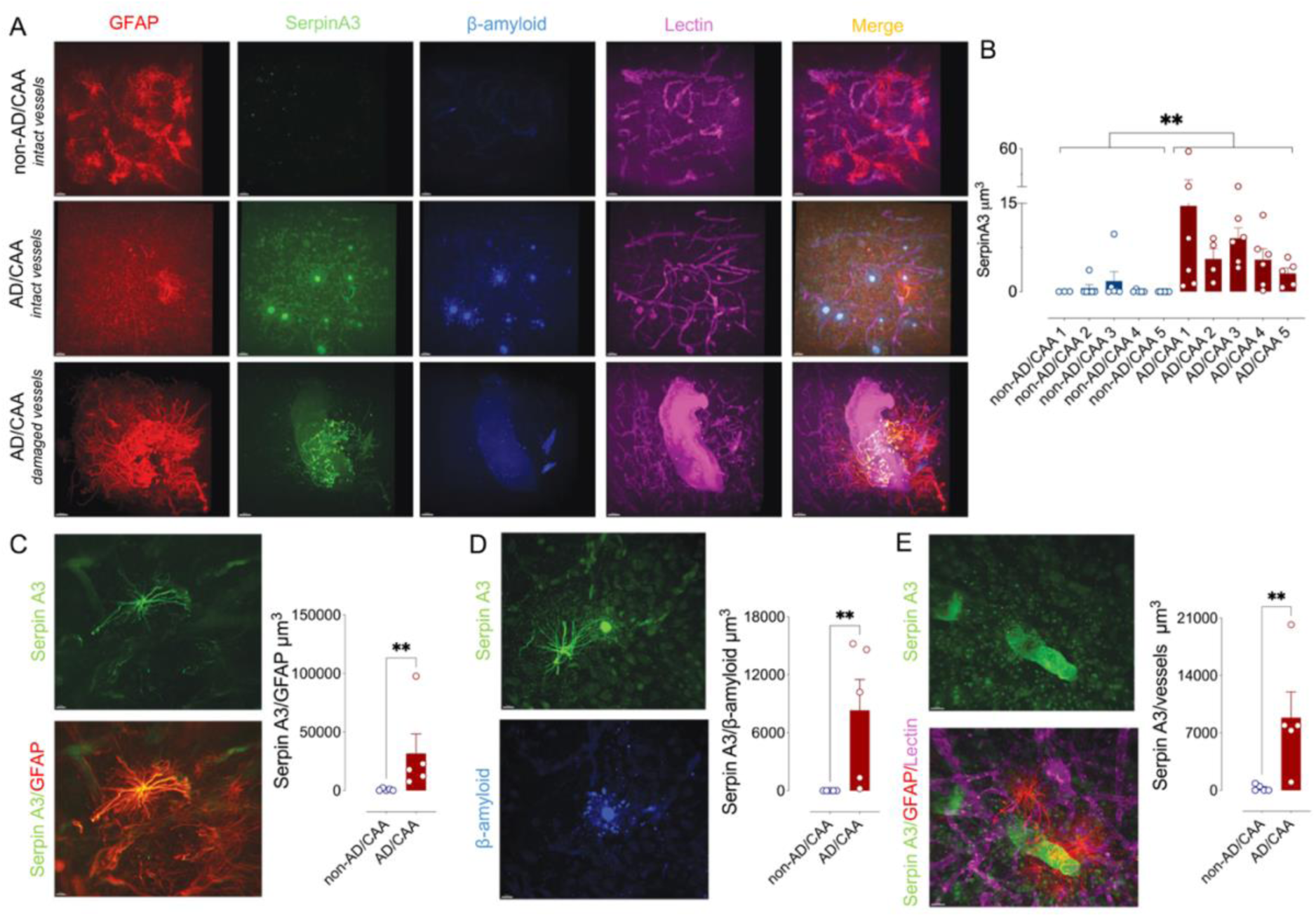
Three-dimensional reconstruction shows Serpin A3 associated with ß-amyloid, GFAP-positive astrocytes, and the vasculature in cerebral amyloid angiopathy. A) Representative three-dimensional light-sheet image of Serpin A3 (green) in human tissue across distinct vascular pathological conditions. Non-AD/CAA vessels (top panel), AD/CAA vessels (middle panel), scale bar 50 µm, and AD/CAA structurally compromised vessels (bottom panel), scale bar 100 µm. Astrocytes were labeled with GFAP (red), while vessels were labeled with Tomato-Lectin (magenta); β-amyloid deposits, neuritic plaques, neurofibrillary tangles, and other tau aggregates were visualized with Methoxy-04 (MX-04, blue). B) Quantification of Serpin A3 immunofluorescence in three-dimensional human tissue sections. Bars show all measurements for each donor, illustrating heterogeneity within and between groups. For statistical analysis, measurements were averaged for each donor, and each average was treated as one independent biological replicate (*n*=5 patients per group). Data are presented as mean ± S.E.M. \*\**p*<0.01 by Mann-Whitney U test. C) Representative image of Serpin A3 (green, top panel) and double stained with GFAP (red, bottom panel), scale bar 50 µm. To assess astrocytic Serpin A3 levels, we measured the ratio of Serpin A3 volume within GFAP-positive volumes. Data are represented as mean ± S.E.M., *n=5*donors per group. \*\**p*<0.01 by Mann-Whitney U test. D) Representative image of Serpin A3 (green, top panel) associated with MX-O4-stained β-amyloid (bottom panel), scale bar 50 µm. The graphic shows the ratio of Serpin A3 volume within MX-O4-positive volumes. Data are represented as mean ± S.E.M., *n=5* patients per group. \*\**p*<0.01 by Mann-Whitney U test. E) Representative image of Serpin A3 (green, top panel) on vessels (magenta, also showing GFAP in red, bottom panel), scale bar 50 µm. The graphic shows the ratio of Serpin A3 volume within vascular volumes. Data are represented as mean ± S.E.M., *n=5* patients per group. \*\**p*<0.01 by Mann-Whitney U test.

### Serpin A3 is prominently increased in a case of severe amyloid-related imaging abnormalities

To evaluate the level and pattern of Serpin A3 in the context of severe ARIA, we analyzed tissue from a previously reported fatal case of lecanemab-associated cerebral β-amyloid-related arteritis [25]. Serpin A3 immunoreactivity was prominent around microhemorrhages (Fig. 5A and supplementary video, see Fig. S3) and dilated/degenerating vessels (Fig. 5B). Serpin A3 levels were markedly elevated in the setting of severe ARIA, higher than those observed in the other CAA cases (Fig. 5C). Serpin A3 in this case was detected in astrocytes and also in non-astrocytic cells.

**Figure 5.**
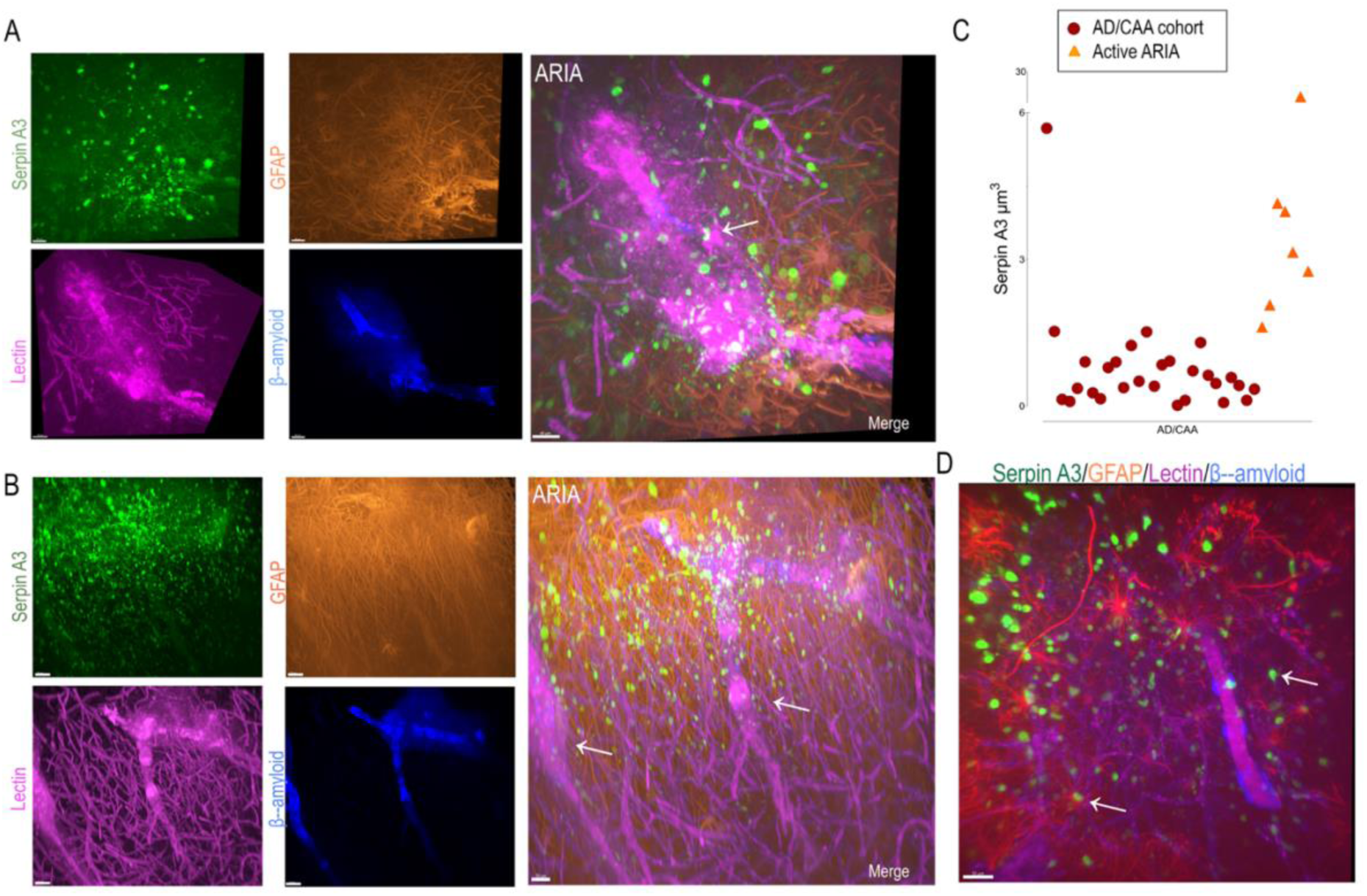
Serpin A3 is increased in amyloid-related imaging abnormalities (ARIA). A) Three-dimensional light-sheet microscopy of a case of active, severe ARIA immunostained for Serpin A3 (green), astrocytes (GFAP, red), vasculature (Tomato-Lectin, magenta), and β-amyloid (Methoxy-04, blue). The merged panel shows a focal microbleed (arrow), with Serpin A3 immunoreactivity localized adjacent to the affected vessel, including some astrocytes Serpin A3 positive, Scale bar, 50 μm. For more details, see also the supplementary video. B) The same case showing a region of vascular dilation (arrow), with Serpin A3, GFAP, Tomato-Lectin, and β-amyloid immunoreactivity as in (A). Scale bar, 50 μm. C) Scatter plots illustrate the comparison of Serpin A3 volume (μm^3^) between AD/CAA cases and an individual pathological ARIA case (red*; n*=28 fields from 5 AD/CAA donors and orange triangles; *n*=7 fields from a severe lecanemab-associated ARIA case. D) Another example image from the same ARIA case, showing Serpin A3 astrocytes positive (arrow) and Serpin A3 positivity in non-astrocytic cells (arrow). Scale bar, 50 μm.

## Discussion

Our understanding of the heterogeneity of astrocytes and the breadth of biological roles they play in the brain is rapidly increasing. Astrocytes are perhaps best known for their roles in supporting neuronal physiology, but they also make up a critical component of the neurovascular unit. Their abundance in the brain closely parallels endothelial populations. Astrocytes have been classically separated into homeostatic and activated states; however, transcriptomic and spatial profiling studies have revealed diverse, and molecularly distinct, astrocyte populations associated with specialized anatomical niches and functional roles [26]. Among these, Serpin A3-positive astrocytes have emerged as an inflammatory and tissue injury-responsive subtype[27–29]. Our findings extend this framework by identifying an association between Serpin A3 and cerebrovascular degeneration in CAA and ARIA).

We found that Serpin A3 protein abundance was increased in brain tissue from patients with AD and CAA. The pattern of increased Serpin A3 is heterogeneous and is associated with areas of tissue pathology. We observed Serpin A3 around some β-amyloid plaques, confirming a prior report [30]. Additionally, we observed Serpin A3 to be associated with the vasculature in CAA, particularly in astrocytes surrounding arterioles with degenerative morphologies, including microaneurysmal dilation [31]. Because of the association of CAA with intracerebral hemorrhage, the preferential accumulation of Serpin A3 within and around degenerating rather than structurally intact vessels links Serpin A3 could suggest a link to cerebrovascular fragility.

Given its function as a serine protease inhibitor, Serpin A3 may influence extracellular matrix modifying enzymes and vascular stability by modulating protease-mediated connective tissue remodeling. In ARIA, which is an accelerated form of CAA induced by anti-amyloid immunotherapy [25], Serpin A3 was strongly associated with inflamed vessels, with expression in astrocytes and non-astrocytic cells. Given the extent and acuity of vascular remodeling in this condition, a mechanistic role for Serpin A3 in ARIA should be further explored.

In contrast, Serpin A3 mRNA tended to be reduced in AD brains, especially within the vascular compartment. Discrepancies between transcriptomic and protein abundance findings are common among secreted proteins [32], likely reflecting differences in extracellular accumulation, stability, and clearance. Our results indicate that increased Serpin A3 levels occur within a small, reactive astrocyte subpopulation associated with degenerating vasculature rather than through widespread mRNA expression across the tissue. Transcriptomic analyses frequently underrepresent vascular and perivascular compartments, so the snRNAseq transcriptomic result should be interpreted as reflecting parenchymal expression levels. Additionally, the total level of expression of Serpin A3 was quite low, with a significant percentage of cells not expressing any detectable Serpin A3. Oligodendrocyte expression of Serpin A3 has been previously reported, especially in the setting of stroke [33]. We observed oligodendroglial Serpin A3 expression in the snRNAseq database and oligodendrocyte precursor cell expression of Serpin A3 correlated with longitudinal cognitive. We do also note that while some of the elevation in Serpin A3 in the ARIA case was associated with astrocytes, a substantial portion was in non-astrocytic cells, so oligodendroglial Serpin A3 is possible in ARIA. Finally, a few writers have proposed that Serpin A3 from the periphery may be the source or a source of Serpin A3/Serpin A3n in the brain, and we cannot exclude this possibility [9]. Nevertheless, on balance it appears that astrocytes are the major source of Serpin A3 in the brain in CAA, especially in the perivascular niche.

Serpin A3 is a serine protease inhibitor best known for binding and inhibiting cathepsin G, although its substrates are diverse. Previous *ex vivo* studies suggested that Serpin A3 may regulate BBB function, based on a modest reduction in Claudin 5 level and modest increase in VCAM1 level in murine brain explants cultured in high concentrations of exogenous Serpin A3n, the murine homolog of Serpin A3 [6]. The primary vascular site of Serpin A3 detection in the brain in this study was arterioles, not capillaries (although some capillary staining was observed in the hippocampus). This may argue against a major role for Serpin A3 in BBB function, although conceivably it could regulate aspects of the BBB in a regionally specific manner in sites with capillary expression. Increased Serpin A3 expression in AD has been reported in the human parahippocampal gyrus (see Agora AD Knowlege Portal: agora.adknowledgeportal.org/genes/ENSG00000196136/evidence/rna), which could account for its increased presence in the local neurovascular unit in the adjacent hippocampus. In the ROSMAP dataset, higher levels of astrocytic Serpin A3 correlated with better longitudinal cognitive performance, which is not, on its face, consistent with the idea that Serpin A3 impairs critical functions in the brain at physiologic levels. Instead, the preferential association of Serpin A3 with degenerating arterioles is more suggestive of a role in structural remodeling of the vessel and/or a response to vascular injury. This demonstrates the importance of interpreting *in vitro* models within the anatomical and pathological context of relevant human tissues.

The upstream regulatory pathway for Serpin A3 is thought to involve TNF-α and STAT3 signaling [6]. Both STAT3 and Serpin A3 are upregulated in astrocytes carrying AD-associated presenilin mutations [22]. Growing evidence indicates that APOE-associated inflammatory astrocyte programs converge on overlapping pathways regulating reactive gliosis and cerebrovascular dysfunction. In fact, recent work demonstrated that Serpin A3n interacts with APOE, and Serpin A3n knockdown resulted in APOE downregulation [34]. Together, these findings support the possibility that Serpin A3 participates in broader APOE- and STAT3 associated astrocytic inflammatory responses contributing to vascular remodeling and CAA progression.

Although Serpin A3n is frequently described as the murine counterpart of human Serpin A3, species-specific differences should be considered. Mice possess an expanded Serpin 3n paralog cluster with potential differences in Serpin A3/Serpin A3n substrate specificity, regulation and disease context. Recent work has revealed broad, but regionally heterogeneous distribution of Serpin A3n in the mouse CNS, with enrichment in the striatum, cerebellum, and thalamus while the highest astrocytic abundance was found in the hypothalamus [26]. The primary molecular target of human Serpin A3 is cathepsin G, whereas murine Serpin A3n predominantly binds granzyme B and leukocyte elastase (in addition to cathepsin G), the latter of which in particular may partly mediate BBB effects in mice. The increased amount of Serpin A3 within degenerating vessels suggests that Serpin A3 may actively contribute to pathological extracellular remodeling and impaired vascular homeostasis during CAA progression rather than simply reflecting reactive astrocyte activation.

There is some evidence that Serpin A3 is expressed in non-astrocytic cell types. Our results using snRNAseq data from the ROSMAP study support this view, showing that vascular fibroblasts and endothelial cells also express considerable levels of Serpin A3, while neurons, oligodendrocytes and microglial express lower levels. Serpin A3 is also expressed in neural precursor cells where it likely plays a role in brain development. While Serpin A3n knock-out mice are neurologically normal, there is some evidence that Serpin A3 knock-in mice can induce gyrification in the mouse brain and improve memory, although not all reports have supported this conclusion [35].

Finally, Serpin A3 acts as one of several enzymes critical to remodeling vessels that are likely modulated together in injury responsive astrocytes and almost certainly act in concert.

These pathways need to be understood comprehensively to define the mechanism of vascular injury in neurodegenerative diseases. In this context, increased presence of Serpin A3 around degenerating vessels may be a compensatory response to vascular injury, potentially blocking protease-mediated extracellular matrix degradation and vessel wall destabilization.

## Conclusions

Serpin A3, also called α1-antichymotrypsin, has long been known as a marker of activated, injury-responsive astrocytes. The expression of Serpin A3 is potently induced by inflammatory signaling molecules including STAT3, TNF-α, IL-6 and IL-1β and Serpin A3 is thought to serve a protective role in limiting inflammatory tissue injury. It has been linked to Alzheimer’s disease, multiple sclerosis, Parkinson’s disease, brain trauma and amyotrophic lateral sclerosis. Our study demonstrates that Serpin A3 is increased in CAA and AD and linked to degenerating vessels.

## Acknowledgements

We thank the many patients and their families who have donated brain tissue to support this and other necessary studies. We also acknowledge the contributions of the Vanderbilt Cell Imaging Shared Resource (CISR) Core.

## Author contributions

The authors confirm contribution to the paper as follows: study conception and design: L.V-A., W.R-F. and M.S.S.; snRNAseq analyses: V.A.J and T.J.H; western blot analyses and antibody validation: W.R-F, N.W., C.J.W.; microscopy: L.V-A., W.R-F, S.A.H, B.C.M, H.B, M.S.S; analysis and interpretation of results, draft manuscript preparation, and substantial revision: L.V-A., W.R-F., N.W., C.J. W, S.A.H, V.A.J., H.B, T.J.H, B.C.M and M.S.S. All authors contributed to the final revision of the manuscript and approved the version submitted for publication.

## Funding

This project is financially supported by a grant from the National Institutes of Health (1R01NS129735).

## Data availability statement

The data supporting this study’s findings are available from the corresponding author upon reasonable request.

## Competing interests

MSS has received research support from the NIH (1RF1NS129735, R01AG078803, RF1NS130334, and R21AG070859), American Federation of Aging Research (K76AG060001/Beeson Scholar), the NeuroScience Innovation Foundation. He has received payments for consulting from Guidepoint and has received payments for expert testimony in medicolegal cases. He chaired the DSMB for the REVISION trial, serves on the medical advisory board for Septagen and is the research integrity officer for the Journal of Alzheimer’s Disease. TH serves on the scientific advisory board for Circular Genomics and serves as the Deputy Editor for Alzheimer’s & Dementia: TRCI.

The rest of the authors declare no competing interests.

## Notes

https://doi.org/10.6084/m9.figshare.34672842

