## Supplementary material for "Serpin A3 is Increased in Cerebral Amyloid Angiopathy and ARIA"

**SUPLEMENTATY MATERIAL**

**Supplementary Table 1.** Samples demographics de-identified neuropathological information.

| Age at death / Sex | Primary clinical categorization | AD neuropathology | CAA grade | PMI | Other neuropathologies |
|---|---|---|---|---|---|
| 71/M | Alzheimer's disease | Severe (A3, B3, C3) | Severe CAA | 12h | Right parietal hemorrhage |
| 81/M | Alzheimer's disease | Low (A2, B1, C2) | Severe CAA | 19h | Hemorrhages |
| 81/F | Alzheimer's disease | Severe (A3, B3, C3) | Severe CAA | 35h53' | None |
| 68/F | Alzheimer's disease | Severe (A3, B3, C3) | Severe CAA | 9h | None |
| 68/F | Alzheimer's disease | Severe (A3, B3, C3) | Low CAA | 15h | None |
| 70/M | Alzheimer's disease | Severe (A3, B3, C3) | Severe CAA | 11h | Tau in middle and superior temporal gyrus |
| 61/M | Alzheimer's disease | Moderate-severe (A2, B2, C3) | Moderate-severe CAA | 10h | None |
| 73/M | Alzheimer's disease | Moderate (A3, B0-1, C3) | Severe CAA | 10h | Multiple ICH, CMB |
| 77/M | Alzheimer's disease | Low (A2, B1, C2) | Severe CAA | 11h | Frontal lobe and old temporal ICH |
| 66/F | Alzheimer's disease | Severe (B5, Thal 4, C3) | Mild CAA | 25h | Small subdural |
| 73/M | Alzheimer's disease | Severe (A3, B3, C3) | Moderate CAA | 7h55' | None |
| 82/M | Alzheimer's disease | Low (A3, B0, C2) | Mild CAA | 8h | Hippocampal sclerosis, diffuse Lewey body dementia |
| 65/M | Alzheimer's disease | Severe (A3, B3, C3) | Moderate CAA | 49.5h | Mixed dementia, LBD brainstem |
| 79/M | Alzheimer's disease | Low (A3, B1, C1) | Mild CAA | 7h | None |
| 68/M | Alzheimer's disease | Moderate (A3, B2, C3) | Mild CAA | 18h40' | FTD TDP43, 4R-tau grains |
| 88/F | Alzheimer's disease | Moderate (A3, B2, C2) | Severe CAA | 14h | None |
| 60/F | Alzheimer's disease | Severe (A3, B3, | Mild CAA | 15h | LBD |

| | | | | | |
|---|---|---|---|---|---|
| | | C3) | | | |
| 76/M | Alzheimer’s disease | Severe (A3, B3, C3) | Severe CAA | 14h | None |
| 76/M | Alzheimer’s disease | Severe (A3, B3, C3) | Moderate CAA | 4h | None |
| 76/M | Alzheimer’s disease | Moderate (A3, B2, C2) | Moderate CAA | 21h | ICH |
| 67/F | Alzheimer’s disease | Severe (BRAAK VI) | Focally severe | 28h | None |
| 65/F | Alzheimer’s disease | Severe (BRAAK VI) | Focally severe | 19h | None |
| 92/F | Alzheimer’s disease | Severe (BRAAK VI) | Mild | 16h | None |
| 78/F | Alzheimer’s disease | Severe BRAAK V | Mild | 6h | None |
| 62/M | Cerebral amyloid angiopathy | None | Moderate CAA | 35h | Hemorrhages |
| 76/F | Cerebral amyloid angiopathy | Mild (A3, B1, C1) | Severe CAA | 38h | None |
| 68/M | Cerebral amyloid angiopathy | Mild (A3, B1, C1) | Severe CAA | 100h | None |
| 62/M | Neurological control | None | None | 11h3’ | TIA |
| 68/M | Neurological control | None | None | 17h18’ | None |
| 27/M | Neurological control | None | None | 11h15’ | None |
| 45/M | Neurological control | None | None | 13h1’ | Cerebrovascular atherosclerosis and thrombosis |
| 87/M | Neurological control | None | None | 12h30’ | None |
| 71/M | Neurological control | None | None | 29h5’ | Moderate vascular disease |
| 87/M | Neurological control | None | None | 6h12’ | LMCA stroke |
| 66/F | Neurological control | None | None | 33h12’ | None |
| 72/M | Neurological control | None | None | 12h9’ | None |
| 68/M | Neurological control | None | None | 24h | None |

| | | | | | |
|---|---|---|---|---|---|
| 66/M | Neurological control | None | None | 23h38’ | None |
| 70/F | Neurological control | None | None | 35h15’ | None |
| 29/F | Neurological control | None | None | 26h | Pulmonary hypertension |
| 24/F | Neurological control | None | None | 6h | None |
| 102/F | Neurological control | None | None | 47h | - |
| 83/F | Neurological control | None | None | 6h | - |
| 78/M | Neurological control | None | None | 20h | - |
| 65/F | Neurological control | None | None | 17h | - |

**Fig. S1A.** Rabbit anti-Serpin A3 (GTX55503, GeneTex) validation for western blotting. Increasing amount of purified Serpin A3 was probed by western blot and detected a single, specific immunoreactive band around 50 kDa.

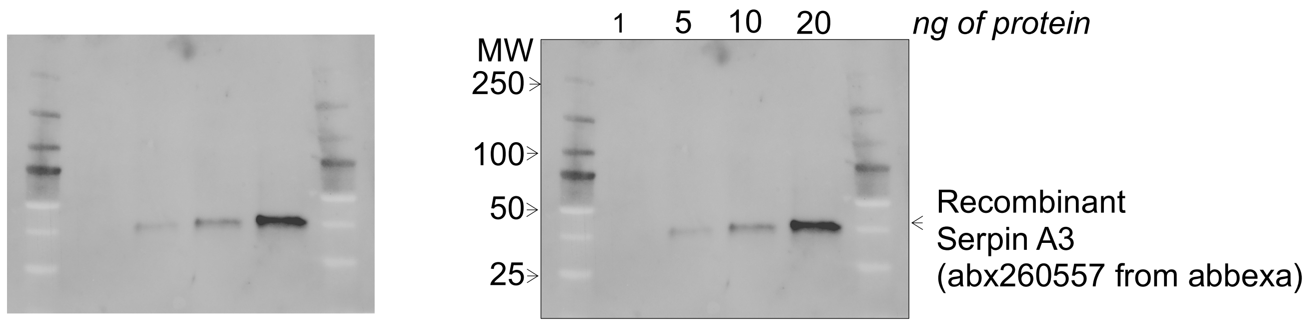


**Fig. S1B.** Rabbit anti-Serpin A3 (GTX55503, GeneTex) validation for immunohistochemistry. HeLa cells expressing recombinant fluorescent-tagged SerpinA3 (green) were immunolabeled with the antibody (red), showing concordant localization. Scale bar, 20 µm.

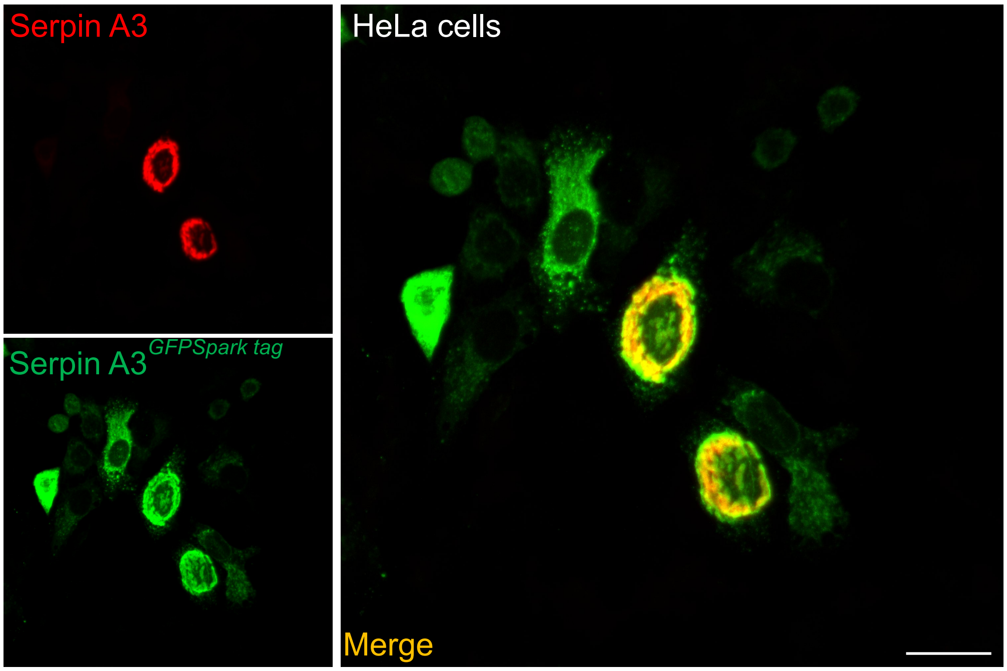

**Fig. S3.** Raw data of Serpin A3 levels in grey matter. Western blot of Serpin A3 and GAPDH in protein fraction from gray matter. Each lane corresponds to a donor. X; excluded samples.

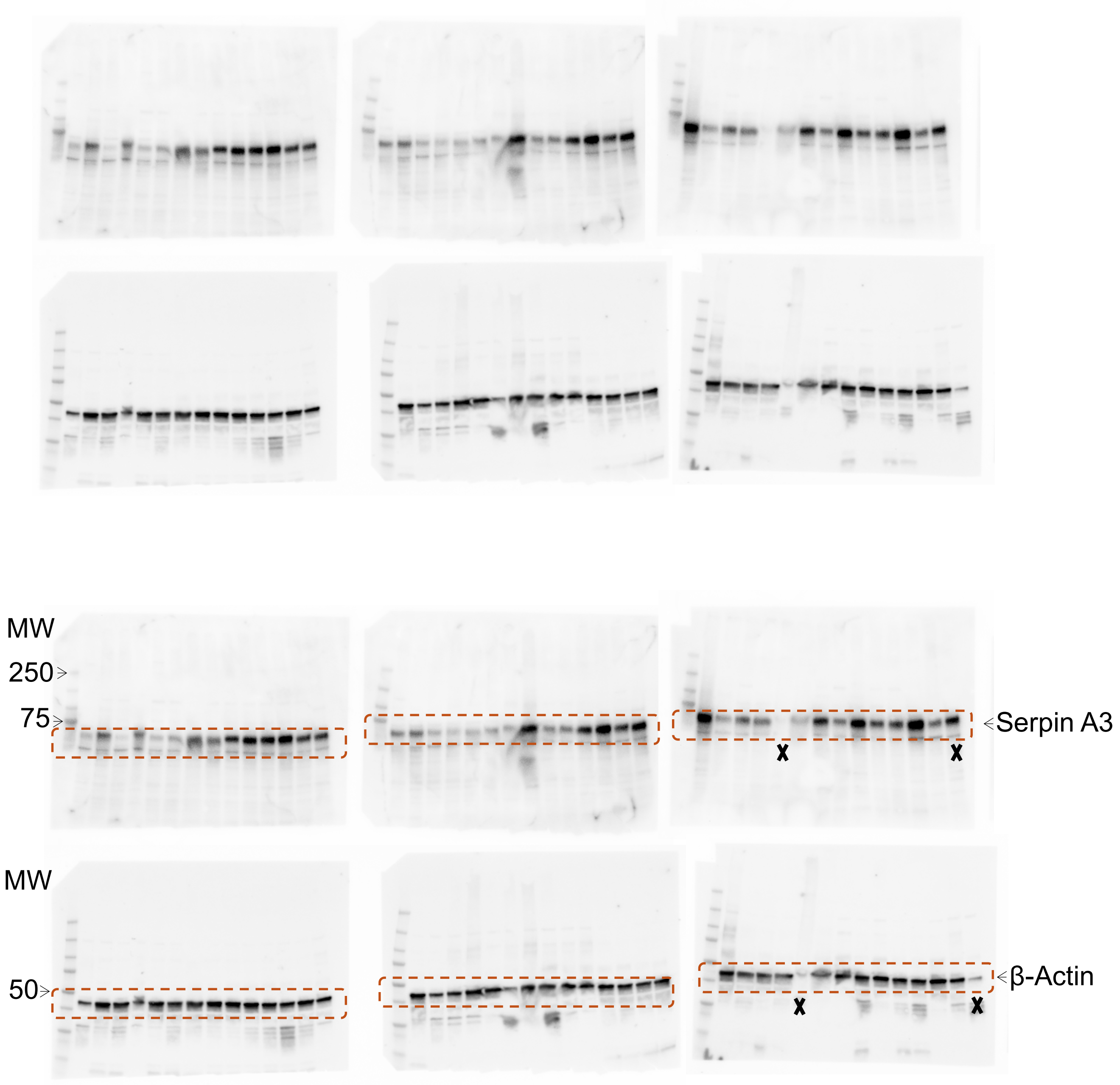

**Fig. S4. Representative video illustrating three-dimensional analysis.** Three-dimensional images acquired using light-sheet microscopy, immunoassayed for Serpin A3 (green), astrocytes (GFAP, red), vasculature (Tomato-Lectin, magenta), and β-amyloid (Methoxy-04, blue). Showing microbleeding vessels.

See, https://doi.org/10.6084/m9.figshare.34672842
